# Continuous Reports of Sensed Hand Position During Sensorimotor Adaptation

**DOI:** 10.1101/2020.04.29.068197

**Authors:** Jonathan S. Tsay, Darius E. Parvin, Richard B. Ivry

## Abstract

Sensorimotor learning entails multiple learning processes, some volitional and explicit, and others automatic and implicit. A new method to isolate implicit adaptation involves the use of a “clamped” visual perturbation in which, during a reaching movement, visual feedback is limited to a cursor that follows an invariant trajectory, offset from the target by a fixed angle. Despite full awareness that the cursor movement is not contingent on their behavior, as well as explicit instructions to ignore the cursor, systematic changes in motor behavior are observed, and these changes have the signatures of implicit adaptation observed in studies using classic visuomotor perturbations. While it is clear that the response to clamped feedback occurs automatically, it remains unknown if the adjustments in behavior remain outside the participant’s awareness. To address this question, we used the clamp method and directly probed awareness by asking participants to report their hand position after each reach. As expected, we observed robust deviations in hand angle away from the target (average of ∼18°). The hand reports also showed systematic deviations over the course of adaptation, initially attracted towards the visual feedback and then in the opposite direction, paralleling the shift in hand position. However, these effects were subtle (∼2° at asymptote), with the hand reports dominated by a feedforward signal associated with the motor intent yet modulated in a limited way by feedback sources. These results confirm that adaptation in response to a visual perturbation is not only automatic, but also largely implicit.

**NEWS AND NOTEWORTHY:** Sensorimotor adaptation operates in an obligatory manner. Qualitatively, subjective reports obtained after adaptation demonstrate that, in many conditions, participants are unaware of significant changes in behavior. In the present study, we quantified participants’ awareness of adaptation by obtaining reports of hand position on a trial-by-trial basis. The results confirm that participants are largely unaware of adaptation, but also reveal the subtle influence of feedback on their subjective experience.

## INTRODUCTION

Motor adaptation is the process of calibrating well-practiced actions to maintain performance in response to changes in the environment or body. A large body of work has focused on how sensory prediction error, the difference between predicted and actual sensory feedback, drives motor adaptation in an automatic manner (Shadmehr, Smith, & Krakauer, 2010). For instance, if a fatigued ping-pong player begins to produce shots that land close to the net instead of the opponent’s back line, her motor commands would be re-calibrated to result in more forceful movements.

Perturbations of the visual feedback have offered one approach to study motor adaptation in the laboratory. In visuomotor rotation tasks, participants are initially trained to reach to visually defined targets, with veridical feedback of their hand position represented by a cursor. Following this baseline period, a rotation is imposed between the position of the hand and the position of the cursor. To counteract the rotation, the motor system must adjust future movements, generating commands that lead to hand movements in the opposite direction of the perturbation.

While participant’s phenomenological experience *after* learning suggest that the change in behavior is largely implicit (at least for rotations up to 45°), recent methods using probes continuously *during* learning (e.g., aim reports) have made clear that standard visuomotor rotation tasks elicit multiple learning processes (Bond & Taylor, 2015; Mazzoni & Krakauer, 2006; Shmuelof et al., 2012; Taylor & Ivry, 2011; Taylor, Krakauer, & Ivry, 2014). These standard visuomotor tasks conflate sensory prediction errors with task performance errors: The former is assumed to be the driving force for implicit adaptation, whereas the latter has been shown to elicit more strategic changes in performance (Taylor et al., 2014; Werner et al., 2015). Thus, explicit changes in action selection operate in parallel with implicit changes occurring within the motor execution system.

To study sensorimotor adaptation in the absence of strategy use, Morehead et al. (2017) introduced a “visual error clamp” method. As with standard visuomotor rotation tasks, participants reach to a visual target, with feedback limited to a cursor that is time-locked to the radial distance of the hand from the starting position. However, with the clamp method, the cursor follows an invariant path, always offset from the target by a fixed angle. Thus, unlike standard adaptation tasks, the angular position of the feedback is ***not*** contingent on the participant’s behavior. Despite being fully informed of the manipulation and instructed to always reach directly to the target, the participant’s behavior exhibit all of the hallmarks of implicit adaptation, with the heading angle gradually shifting in the direction opposite the clamped feedback. Presumably, this change is driven because the adaptation system, in an obligatory manner, treats the discrepancy between the target and feedback cursor as a sensory prediction error. Since the “error” never changes, the learning function can be observed in the absence of other sources concerning performance (e.g., the reduction in task error that occurs in standard adaptation tasks). Quite strikingly, the change in heading angle will continue for a few hundred trials, reaching asymptotic values that average over 20°, even reaching values greater than 45° in some participants (Kim, Morehead, Parvin, Moazzezi, & Ivry, 2018).

If the visual error clamp indeed elicits implicit motor adaptation, we should expect that adaptation proceeds not only automatically but also unconsciously. The presence of a persistent aftereffect once the clamped perturbation is removed indicates that participants are unaware of their (often substantial) adaptive changes. This behavior is in accord with the participants’ subjective reports: when queried at the end of the experimental session, participants generally report that they had followed the instructions, reaching directly to the target throughout the experiment.

Here we took an alternative tack to these indirect or retrospective probes on awareness, assaying participants’ awareness of the ongoing changes in behavior over the course of adaptation. Specifically, we asked the participant to report the position of their hand *after* each reach during the visual error clamp manipulation. If participants are unaware of their adapted behavior, then the reported hand positions should be at the target location, with some variation due to motor and perceptual noise. Alternatively, participants may respond to the clamped error in an implicit and obligatory manner but also be aware of the resulting change in behavior. In the extreme, the hand reports would track the true hand position. Such an outcome would be reminiscent of the alien hand sign (Brion & Jedynak, 1972), a condition in which patients are aware that they are producing “unintended” movements, but cannot volitionally control these movements.

## METHODS

Young adults (n = 32, 21 female, mean age = 21, age range: 18 - 25) were recruited from the Berkeley community. All participants were right-handed, as verified with the Edinburgh Handedness Inventory (Oldfield, 1971). Participants received course credit or financial compensation for their participation. No statistical methods were used to determine the target sample sizes; rather, the sample sizes were based on previous studies using the error clamp method (Kim et al., 2018; Morehead et al., 2017; Parvin, McDougle, Taylor, & Ivry, 2018). The protocol was approved by the institutional review board at the University of California, Berkeley.

### Reaching Task

Participants were seated at a custom-made table (Fig. 1a) that housed an LCD screen (53.2 cm by 30 cm, ASUS) mounted 27 cm above a digitizing tablet (49.3 cm by 32.7 cm, Intuos 4XL; Wacom, Vancouver, WA). The participant made reaching movements by sliding a modified air hockey “paddle” that contained an embedded stylus. The tablet recorded the position of the stylus at 200 Hz. The experimental software was custom written in Matlab, using the Psychtoolbox extension (Brainard, 1997).

**Fig. 1.**
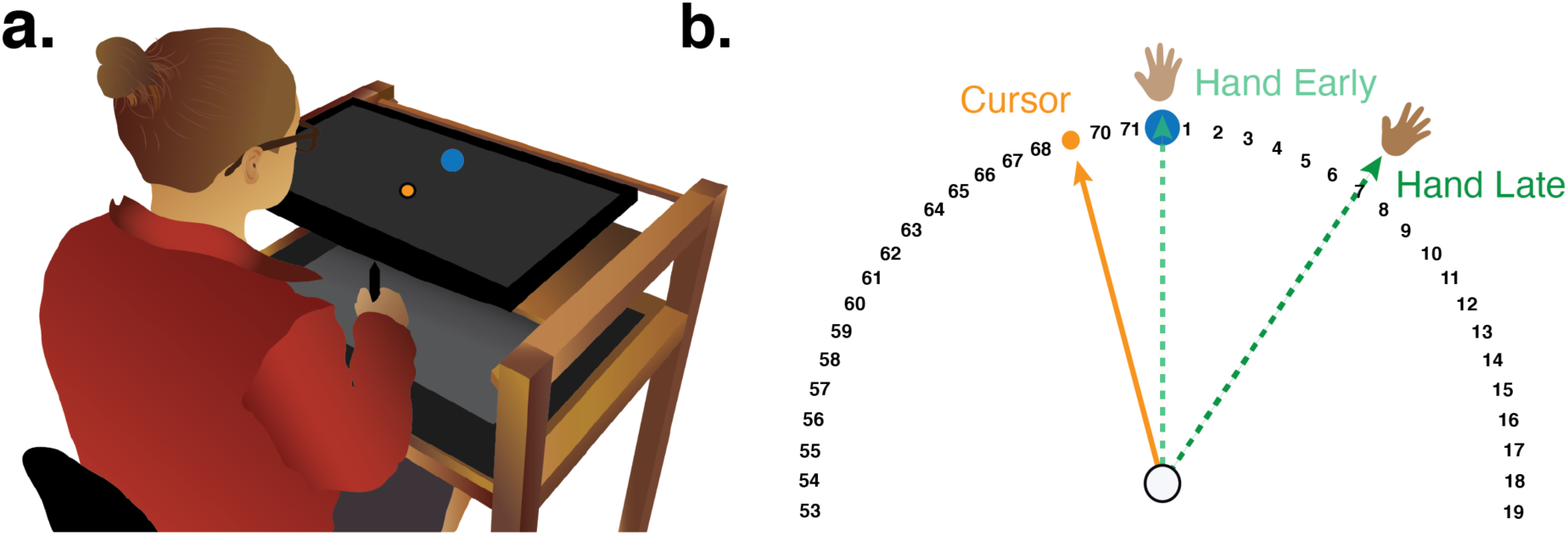
Experimental Methods. **(a)** Experimental apparatus and setup. **(b)** Schematic overview of the error clamp paradigm, in which the angular path of the cursor (yellow) is held constant, and independent of hand movement direction (green). Dotted lines depict representative trajectories at the start (Early) and end (Late) of the error clamp block.

On each trial, the participants made a center-out planar reaching movement from the center of the workspace to a visual target. The center position was indicated by a white circle (0.6 cm in diameter) and the target location was indicated by a blue circle (also 0.6 cm). The target could appear at one of four locations on an invisible virtual circle (45°, 135°, 225°, 315°), with a radial distance of 8 cm from the start location. The monitor occluded direct vision of the hand, and peripheral vision of the arm was minimized since the room lights were extinguished during the experimental session.

To initiate each trial, the participant moved the stylus into the start location. Feedback of the position of the hand, given in the form of a white cursor (0.35 cm diameter), was only provided when the stylus was within 2 cm of the center of the start circle. Once the participant moved the stylus into the start circle and maintained that position for 500 ms, the target appeared. The location of the target was selected in a pseudo-randomized manner, with each location sampled once every four trials. The participant was instructed to reach, attempting to rapidly “slice” through the target. The feedback cursor, when presented (see below) remained visible throughout the duration of the reach and remained fixed for 500 ms at the endpoint location when the movement amplitude reached 8 cm. If the movement was not completed within 300 ms, the message “too slow” was played over the speaker.

The feedback could take one of three forms: veridical feedback, no-feedback, and error clamp feedback. During veridical feedback trials, the location of the visual feedback was veridical, corresponding to the location of the stylus/hand. During no-feedback trials, the feedback cursor was extinguished as soon as the stylus left the start circle and remained off for the entire reach. The cursor only became visible during the return phase of the trial when the stylus was within 2 cm of the start circle. During error clamp trials (Fig. 1b), the cursor moved along a fixed trajectory relative to the position of the target. The clamp was temporally contingent on the participant’s movement, matching the radial distance of the stylus from the center circle (up to 8 cm), but non-contingent on the movement in terms of its angular offset. The fixed angular offset (with respect to the target) was 15° in Experiment 1 and 45° in Experiment 2. The participant was instructed to “ignore the visual feedback and reach directly to the target.”

On some trials, the participants were required to provide a hand report. For these trials, the participant was instructed to maintain their hand position at the end of the outbound segment. A series of numbers appeared as soon as the amplitude of the movement exceeded 8 cm, separated by 5° to form a virtual ring at a radial distance of 8 cm. The numbers (“0” to “71”) ascended in the clockwise direction, with the number “0” positioned at the target location. The participant reported their hand position by verbally indicating the number closest to the perceived location of the stylus.

### Experiment 1

To probe awareness of the consequences of implicit adaptation, the participants (n=16) in Experiment 1 were asked to report their hand position after each reaching movement. The experiment was organized into six blocks of trials. The first three blocks assessed baseline performance in the absence of a perturbation. The first block was composed of 20 reach-only trials without feedback to familiarize the participants with the apparatus. After this block, the hand report procedure was introduced and was included in the remaining five blocks (trials 21 — 360). These consisted of 40 trials with veridical feedback, 40 more trials without feedback, 200 trials with error clamp feedback, 40 trials with no feedback (aftereffect), and a final set of 20 trials with veridical feedback (washout). During the error clamp block, the cursor always followed an invariant trajectory, displaced from the target by 15°. The direction of this displacement was either clockwise or counterclockwise, counterbalanced across participants. Note that we sandwiched the error clamp block with no feedback blocks to provide a measure of adaptation that accounts for idiosyncratic biases in reaching.

Before the error clamp block, the experimenter provided instructions describing the error clamp, emphasizing that its angular trajectory was independent of the participant’s movement and thus, should be ignored. To reinforce the uncoupling of the movement and feedback, three demonstration trials were presented. On the first trial, a target appeared at the 90° location (straight ahead), and the experimenter instructed the participant to first “Reach straight to the left” (i.e., 180°). During the reach, the cursor moved along a trajectory displaced 15° away from the target (matching the direction to be employed with that participant). For the second and third demonstration trials, the target again appeared at 90°, and the participant was instructed to “Reach straight to the right” (0°) and “Reach backward towards your torso” (270°), respectively. For these trials, the cursor again followed a trajectory displaced 15° from the target. After confirming that the participant understood the nature of the feedback, the experimenter again emphasized that the participant should always reach directly to the target and ignore the feedback. The participant then completed the 200-trial block with clamped feedback. Before the 40-trial aftereffect block, the participant was told that no feedback would be provided and that they should continue reaching directly to the target. Prior to the final washout block, the participant was told that the feedback would now correspond to the position of the stylus, and again instructed to reach directly to the target.

### Experiment 2

We repeated the basic hand task in Experiment 2 with a few notable changes. The size of the error clamp was increased to 45° to increase the spacing between the target and the terminal position of the cursor on clamped feedback trials. This manipulation was included to minimize the possibility that, in making their post-reach reports, the participant might confuse the positions of the target and cursor, potentially biasing their reports.

Most importantly, a second error clamp block was added immediately after the first error clamp block in which the direction of the clamp was reversed: If the first clamp block involved a clockwise rotation, the second clamp block involved a counterclockwise rotation, and vice-versa. We expected the hand angle direction would reverse in response to the new clamp, eventually leading to movements in the opposite direction of the reversed clamp. In this manner, we expected to greatly increase the range of changes in hand angle over the course of the experiment. We could exploit this increased range in hand angle to probe whether the hand reports also demonstrate a reversal in direction and increase in range.

Each participant completed 6 blocks: No feedback baseline (20 trials), veridical feedback with hand report (40 trials), no feedback with hand report (40 trials), initial error clamp (180 trials), reversed error clamp (260 trials), and a final washout block with veridical feedback (20 trials). Based on the results of Experiment 1, we reduced the number of trials in the first clamp block to 180, anticipating that participants would be near asymptotic performance. The number of trials in the second clamp block was extended to 260 trials to allow the reversed clamp to first bring the hand angle back towards the target and then reach asymptotic performance in the opposite direction. In this manner, we expected to maximize the range of hand angles, (i.e., essentially double the range over Experiment 1). We did not include a no-feedback aftereffect block given that the results of Experiment 1 showed that the relationship between hand position and hand reports was maintained when the clamped feedback was removed. We opted to conclude with the session with a feedback washout block to ensure that participant’s hand reports remained consistent with their awareness of hand position (i.e., overlapping hand report and hand angle functions).

The hand report procedure lengthens the interval between successive trials. Given that the magnitude of implicit adaptation is impacted by the inter-trial interval (Gary Sing, Bijan Najafi, Adenike Adewuyi, Maurice Smith, 2009) and our desire to maximize the range of hand angles, we opted to use an intermittent procedure to sample the hand reports. These were collected in a set of six report blocks, interspersed across the different reach blocks of the experiment. Hand reports were obtained on trials 21 - 120, 181 - 200, 261 - 300, 361 - 380, 441 - 460, and 521 - 560.

Finally, we modified the procedure used to demonstrate the lack of contingency between the direction of the hand movement and trajectory of the feedback cursor. For the three demonstration trials presented just before the first error clamp block, the target always appeared at the 180° target, and the participant was told to “Reach straight for the target”. Across trials, the feedback cursor terminated at 90° (first trial), 270° (second trial), and 0° (third trial) locations. Following the last demonstration trial, verbal confirmation was obtained that the participant understood that the direction of the cursor was not under his or her control. The experimenter then informed the participant that the cursor feedback would now move in an invariant direction and reinforced the instructions that the participant should ignore the cursor.

There was a mandatory one-minute break between the first error clamp block and the reverse error clamp block. During this break, the experimenter informed participants that the cursor feedback will now follow an invariant trajectory in the opposite direction. Before proceeding, the experimenter obtained verbal confirmation that the participant again understood that the cursor feedback was not tied to his or her movement and should be ignored in its entirety. The participant then completed the 260-trial block with the reverse clamped feedback. Before the last washout block, the experimenter reminded participants to continue reaching directly to the target, with feedback reflecting his or her hand position in a veridical manner.

### Baseline subtraction

The primary dependent variable of reach performance was the hand angle relative to the target, measured at the peak velocity. Outlier responses defined as trials in which the hand angle was greater than 90° from the target location. These were removed from the analysis and constituted only 8 trials out of a total set of 5760 trials.

The hand angle data were pooled over a movement cycle, defined as four consecutive reaches, one to each of the four targets. For each cycle, the means were baseline corrected on an individual basis to account for idiosyncratic angular biases in reaching to the four target locations. These biases were estimated based on heading angles during the last three no feedback baseline blocks (Experiments 1 and 2: cycles 23 – 25), with these bias measures then subtracted from the data for each cycle. For visualization purposes, the hand angles were flipped for blocks in which the clamp was counterclockwise with respect to the target.

The hand report data were converted into angular values, although we note that the reports involve categorical data (numbers spaced at 5° intervals), whereas in angular form they suggest a continuous variable. The hand report data were also baseline corrected on an individual basis to account for idiosyncratic report biases to the four target locations in the exact manner the hand angle data were pre-processed.

### Cluster Permutation Analysis

To evaluate whether participants in Experiment 1 systematically adapted to the visual error clamp, we used a cluster permutation analysis that consisted of two steps. First, a paired t-test was performed for each cycle (after the baseline blocks), asking if the observed hand angle diverged from the hand angle during baseline reaches (cycles 6 to 25). Clusters were defined as epochs of two or more cycles in which *t-*values exceeded a threshold of a *p-*value less than 0.05. The *t-*values were then summed within each cluster to obtain a cluster *t-*score. Second, we compared the observed *t-*scores to the distribution of the maximum absolute *t-*scores (a control for multiple comparisons to limit type-I error rates (Nichols & Holmes, 2002)) obtained from a permutation distribution, which was created by randomly assigning condition labels (baseline or observed hand angle) 1000 times. A *p-*value is obtained by evaluating the proportion of random permutations with *t*-scores greater than the *t-*score from step 1.

The cluster permutation analysis was also used for two analyses relevant to the hand report data. First, a cluster analysis was used to evaluate whether participants’ hand reports during the clamp block significantly deviated from baseline hand reports. Second, a cluster analysis was used to evaluate whether the hand reports significantly deviated from the actual hand angles during the error clamp and aftereffect blocks.

For Experiment 2, we applied the same cluster permutation analysis to evaluate whether the hand angle data for each cycle deviated from baseline (cycles 6 – 25). However, the cluster analysis was not possible for the hand report data because, unlike Experiment 1, these were only obtained intermittently in Experiment 2, violating the cluster test assumption of continuity (Maris & Oostenveld, 2007). Thus, we opted to use two-tailed paired t-tests to compare hand reports during error clamp cycles versus baseline reports. Values are reported as mean ± SEM.

### Other measures of hand angle

For measures of hand angle Experiment 1, we report performance at asymptote (“late adaptation”), quantified as the average of the baseline-corrected hand angle data over the last five error clamp cycles (cycles 71 – 75). The inclusion of a no-feedback block in Experiment 1 also allowed us to measure an aftereffect, defined as the baseline-corrected hand angle of the first cycle from this block (cycle 76).

Similar hand angle measures are reported in Experiment 2. Late adaptation was the average of the baseline-corrected hand angle data over the last five cycles of the first error clamp block (cycles 66 – 70) and the last five cycles of the reverse error clamp block (cycles 131 – 135). We also obtained a range measure by taking the difference between these two measures of late adaptation (cycles 131 – 135 minus cycles 66 – 70).

## RESULTS

### Experiment 1

As expected, participants adapted to the error clamp feedback with the hand angle shifting in the opposite direction of the 15° feedback cursor (Fig. 2). Based on the permutation test, the hand angle deviated from that observed during the baseline block across a large cluster starting from the third cycle of the clamp block (cycles 26 – 75: *t*_*score*_ = 461.41, *p*_*perm*_ *<* 0.001). The mean deviation in hand angle was 17.6° ± 1.7° over the last five cycles of the error clamp block where behavior appeared to be approaching an asymptote. The deviation in hand angle continued to remain substantially higher than baseline throughout the post-clamp aftereffect block in which no feedback was presented (clamp block cycles 76 - 85: *t*_*score*_ = 81.49, *p*_*perm*_ *<* 0.001), providing a second measure of the degree of implicit adaptation. The mean hand angle in this block started close to that observed at the end of the clamp block (15.3° ± 1.5°) and showed a gradual decline over the 10 no-feedback cycles. In summary, we observed robust motor adaptation in response to clamped feedback. Indeed, the response to the clamped feedback was similar to that observed in previous clamp studies (Kim et al., 2018; Morehead et al., 2017), indicating that the hand reports had little, if any impact on adaptation.

**Fig. 2.**
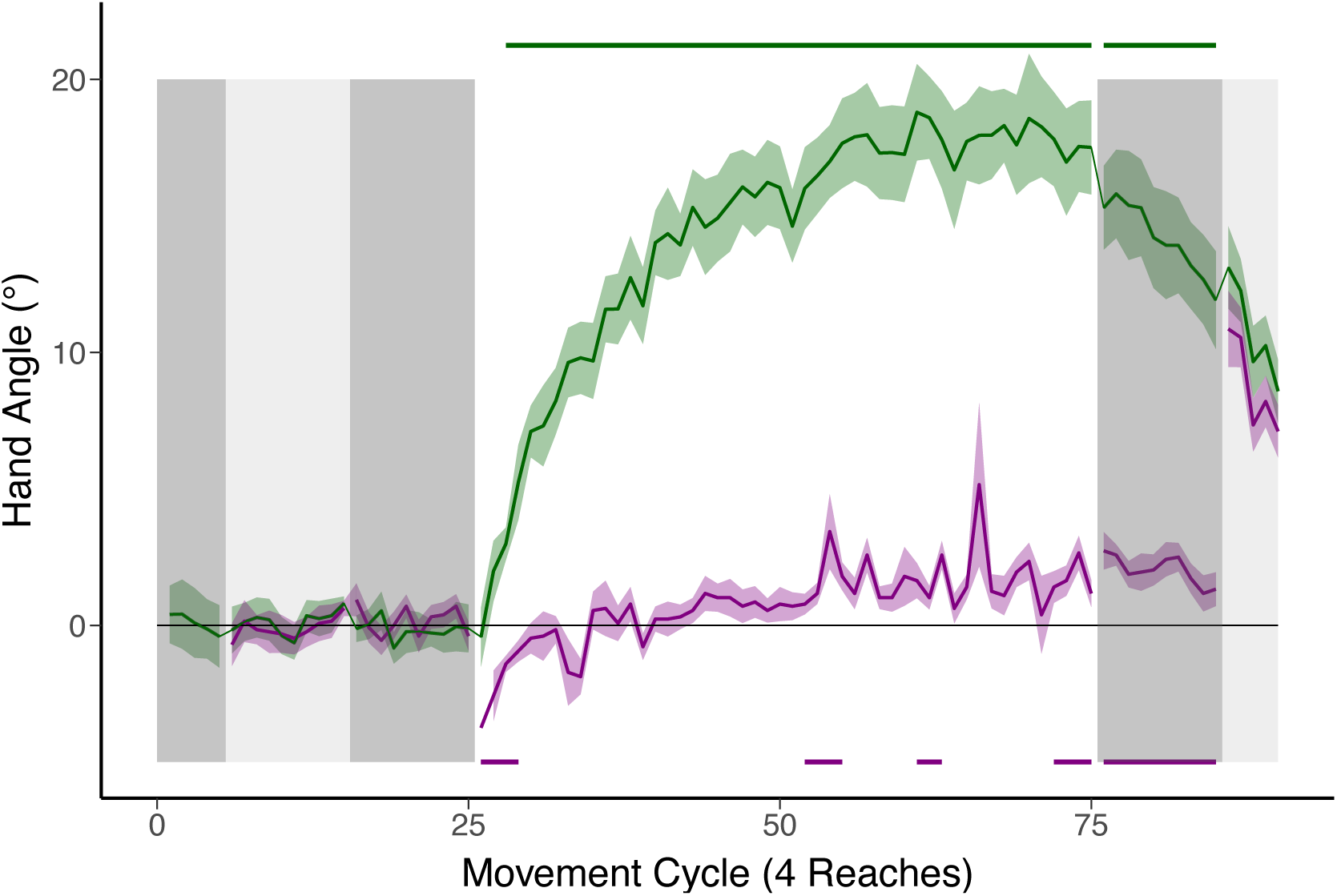
Reaching (green) and hand position report (purple) functions for Experiment 1. Target position is always at 0°. Vertical shading indicates feedback for each block (light gray: veridical; dark grey: no feedback; white: visual error clamp). Clusters in which hand report (bottom, purple) and hand angle (top, green) data are significantly different than baseline are denoted by the bars at the bottom and top of the graph, respectively. SEM denoted by shaded region around each function.

Subjective reports obtained at the end of experiments using a visual clamp indicate that participants are unaware of their adaptation to the visual clamp (Kim et al., 2018; Kim, Parvin, & Ivry, 2019; Morehead et al., 2017). The main goal of this study was to directly probe participants’ awareness of the evolving change in hand angle, asking them to report their hand position after each reach. As can be seen in Figure 2, the hand report data dramatically diverged from the actual hand position, confirming that the observed changes in behavior are largely implicit. To quantify the relationship between the change in hand angle and the participants’ awareness of these changes, we expressed the change in the reported position of the hand as a function of the change in the actual position of the hand. Thus, a large percentage would indicate a close correspondence between the two measures. Focusing on the last five cycles in the clamp block, hand reports account for only 8.3% ± 2.5% of hand angle, revealing little correspondence between the two measures. These data are consistent with the post-report survey data in previous studies, indicating that participants are largely unaware of the large change in motor behavior induced by the error clamp.

However, there are systematic changes in the hand report data during the clamp block. Initially, participants report the hand position to be shifted in the direction of the error clamp, that is, in the opposite direction of the behavioral change (clamp block cycles 26 - 29: *t*_*score*_ = 12.78, *p*_*perm*_ = 0.003, - 2.17 ± 0.62°). Interestingly, this effect was strongest right at the onset of the clamp. One possibility is that some participants were confused by the visual clamp and inferred the position of the hand to be the position of the cursor. This hypothesis would predict that a subset of participants would report hand positions near the clamp location (15°). However, only 9% of all trials in the first block across all participants (22 out of 256 reports) had reports greater than 5° (a conservative cut-off), almost half of which was driven by one participant (9 out of 22 reports). Thus, the shift of perceived hand location towards the clamp suggests that the onset of the visual clamp automatically and implicitly biased the hand reports.

Over time, this initial bias gives way to reports that move in the same direction as the change in hand angle. The reported hand position was reliably different than 0° in the same direction as the actual hand position for only a few clusters (clamp and aftereffect block cycles 53 – 55, 61 – 63, 72 – 75, 71 – 80: all *t*_*score*_ > 9.07, all *p*_*perm*_ *<* 0.03). Even here, the mean values were relatively small (± 2°).

### Experiment 2

Experiment 2 provided a second assay of participants’ explicit experience when adapting to a visual clamp. We introduced a few modifications to the task to focus on two questions. First, we had not anticipated the initial shift in the hand report data in the direction of the clamp. We outlined two hypotheses above: 1) Some participants might have initially interpreted the clamp as veridical feedback or 2) participants may be automatically biased to report their hand position in the direction of the visual clamp. While the hand report data in Experiment 1 support the latter view, we added extra instructions and increased the clamp size from 15° to 45°. Increasing the size of the clamp should reinforce the non-veridical nature of the feedback and thus minimize any possible confusion of the clamp with the hand.

Second, we sought to increase the dynamic range of the change in hand angle, providing a larger window over which to observe changes in the hand reports. We expected the asymptotic change in hand angle (from adaptation) would be largely unchanged in response to the larger clamp angle (Kim et al., 2018). Thus, to increase the dynamic range we employed a design in which the direction of the error clamp was reversed at the midpoint of the experiment. This should result in a shift in the direction of the heading angle for the hand, eventually reaching a similar asymptotic value in the opposite direction. We can then examine if the hand report data shows a similar reversal.

During the initial clamp block, hand angle again deviated in a direction opposite the clamp, the signature of adaptation (Figure 3). The shift in hand was significantly different from baseline by the second error clamp cycle (clamp block cycles 27 – 70: *t*_*score*_ = 341.49, *p*_*perm*_ *<* 0.001). Participants reached an asymptotic value of 18.6 ± 2.7°, similar to the values reported in Experiment 1. When the direction of the clamp was reversed, a corresponding change in hand angle was observed. The mean hand angle crossed the target direction at cycle 85 and reached a maximum (non-asymptotic) mean value of -11.5° ± 2.1°. The deviation in the opposite direction of the clamp was significantly different than the baseline-corrected direction starting at cycle 98 (reversed clamp block cycles 98 – 135: *t*_*score*_ = 203.54, *p*_*perm*_ *<* 0.001). When the effects of the initial and reversed clamp are combined, the summed magnitude of the change in hand angle averaged 30.0° ± 3.9°.

**Fig. 3.**
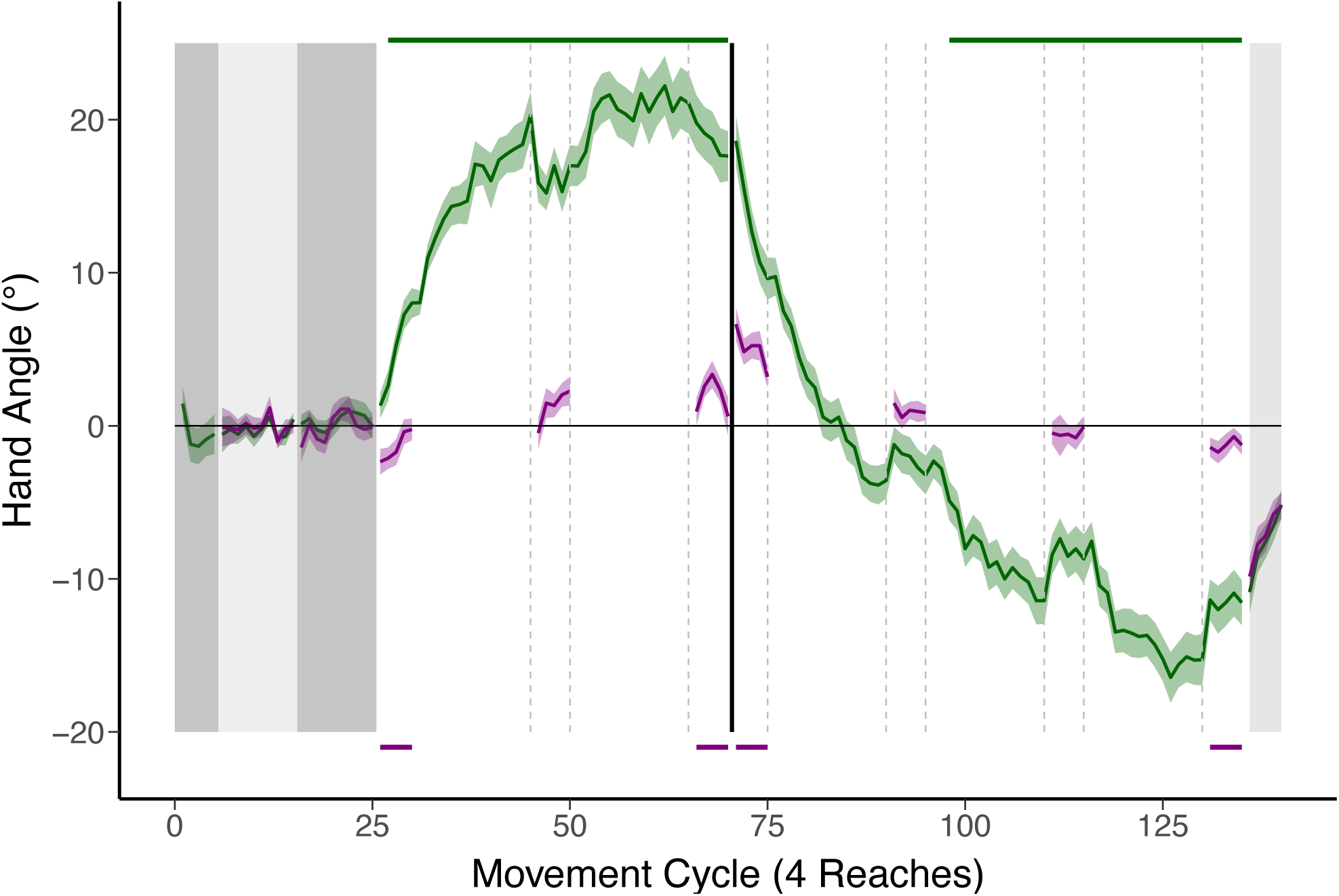
Reaching (green) and hand position report (purple) functions for Experiment 2. Note that hand reports were only obtained in an intermittent manner. Target position is always at 0°. Vertical shading indicates feedback for each block (light gray: veridical; dark grey: no feedback; white: visual error clamp). Black vertical line denotes cycle 71 where the direction of clamped feedback reverses from 45° to - 45°. Clusters in which hand angle deviated from baseline based on permutation test are indicated by green bars at the top of graph. Clusters in which t-test indicated a difference between hand report data and baseline are indicated by purple bars at bottom of the graph. SEM denoted by the shaded region around each function.

We sampled the hand report data in an intermittent fashion in Experiment 2 (purple function in Figure 3). Focusing initially on the subjective reports at the end of each clamp block, we again observed a marked dissociation between the reported and actual position of the hand, confirming that the observed changes in behavior operated largely in an implicit manner. In the last cycles of the first clamp block, the reported change in hand position was only 8.4% ± 5.2% (1.9° ± 3.1°) of the actual change in hand position. A similar dissociation was observed in the reversed clamp block where the hand report positions were 16.3% ± 6.7% (3.2° ± 4.3°) of the actual change in hand angle (with the higher values here due to the fact that adaptation had not reached asymptote in this block).

There were subtle changes in perceived hand position, with a pattern similar to that observed in Experiment 1. Participants again initially perceived their hand position to be shifted in the direction of the clamp, a direction opposite to the evolving change in actual hand position. Given that the hand reports were obtained intermittently, paired t-tests were performed, comparing each mini-block of hand report data to baseline. The shift in the direction of the clamp was significant when averaged over the first mini-block (clamp block cycles 26 – 30: *t*_15_ = −2.93, *p* = 0.01, −1.4 *±* 0.5*°*). The perceived position of the hand then shifted in the direction of the actual hand position, with a reliable difference compared to baseline detected in the third hand report mini-block of the initial clamp block (cycles 66 – 70: *t*_15_ = 2.17, *p* = 0.047, 2.0 *±* 0.8*°*).

When the clamp reversed, we again observed a shift in perceived hand position in the direction of the clamp (cycles 71 – 75: *t*_15_ = 3.57, *p* = 0.003, 3.0 *±* 0.9*°*), that then reversed, following the direction of the actual hand position, becoming reliably different than baseline again in the final hand report mini-block (cycles 131 – 135: *t*_15_ = −2.70, *p* = 0.02, −1.3 *±* 0.5*°*). Importantly, even when reliable, the mean of the hand reports remained near the target at a strikingly small value relative to hand position.

## DISCUSSION

Sensorimotor adaptation is considered an automatic learning process, one that operates in an unconscious manner to ensure that the sensorimotor systems remains calibrated in response to ongoing changes in the state of the body and environmental context (Shadmehr et al., 2010). Several lines of evidence highlight the implicit nature of adaptation. Perhaps most compelling, participants show persistent aftereffects when asked to reach directly to the target during no-feedback blocks, unable to volitionally modify their behavior after being informed that a perturbation is no longer present. Similarly, even when employing a re-aiming strategy to compensate for a large perturbation, a significant portion of the change in heading angle is unaccounted when the participants are asked to report their intended movement direction prior to the reach (Taylor et al., 2014). Indeed, the presence of a sensory prediction error is sufficient to induce adaptation, even when the change in behavior is actually maladaptive in terms of task success (Mazzoni & Krakauer, 2006).

While these observations provide strong support that the behavioral change in adaptation studies occurs in an automatic and implicit manner, it is not clear if participants are aware of the behavioral changes themselves. Probes of awareness obtained at the end of the experiment yield limited information and may be problematic. These retrospective queries are generally framed in a binary manner such as “Did you reach to the target throughout the whole experiment” or “Were you aware of any changes in your hand position”. Moreover, in a standard adaptation study, the task error becomes quite small at the end of the adaptation block, and this reduction in perceived error may impact awareness. Here we assessed awareness in a continuous manner by asking participants to maintain their hand position at the end of the movement and report the angular position of the hand with respect to the target. While this report procedure could be used with standard, contingent visual perturbations, we opted to use the clamp method for two reasons. First, the behavioral change is assumed to be arise from an implicit learning mechanism given that participants are activity discouraged from using an aiming strategy. Second, the perceived “error” remains invariant since the angular direction of the feedback is fixed.

Consistent with the aftereffect data and retrospective reports, the continuous probe of awareness revealed a marked dissociation between the participants’ behavior and their awareness of that behavior: Overall, the clamped feedback elicited a shift in heading angle of ∼18°, yet the participants’ reports of their hand position remained close to the target location, deviating by only ∼2°. Thus, the current results confirm that participants are largely unaware of the behavioral consequences of automatic adaptation.

Nonetheless, the hand report data were not randomly centered about the target as would be expected if participants were oblivious of learning; rather, two systematic changes in the hand reports were observed in both experiments. First, the perceived location of the hand was biased towards the clamped feedback right at the onset of the error clamp block. This effect was similar in response to the introduction of either a 15° or 45° clamp: As such, it seems unlikely to reflect trials in which participants confused the clamped feedback as their veridical hand position. Instead, this initial bias is reminiscent of the proprioceptive shift reported in studies of visuomotor adaptation where the perceived estimate of hand position gravitates towards the visual perturbation (Henriques & Cressman, 2012; Ruttle, Cressman, ‘t Hart, & Henriques, 2016). These proprioceptive shifts have been interpreted as a way in which the brain resolves sensory discrepancies between vision and proprioception to generate a unified estimate of hand position (Ernst & Banks, 2002). This initial bias, observed in both experiments, is consistent both in magnitude (∼4° towards the visual feedback) and rapid onset, with the proprioceptive shift observed in these earlier studies (Cressman & Henriques, 2009, 2010; Ruttle et al., 2016; Salomonczyk, Cressman, & Henriques, 2011).

Second, this bias gave way to a reliable shift in the reported hand position in the direction of adaptation (i.e., away from the visual feedback) that reached a peak of around ∼2°. We assume the reversal in the perceived location of the hand arises from proprioceptive feedback: As adaptation proceeds, veridical feedback from proprioception would signal a hand position that is shifted in the opposite direction of the visual feedback. Consistent with this hypothesis, we observed a positive correlation between the magnitude of adaptation (change in hand angle) and reported hand position at the end of the clamp block (Fig 4). Despite this positive correlation, it is important to keep in mind that there remains a large discrepancy between the actual and reported hand position, with the latter remaining close to the target. The failure to be sensitive to the substantial shifts induced by the clamp may, in part reflect the relatively poor acuity of proprioception, at least when probed in a static manner (Jones, Cressman, & Henriques, 2010).

**Fig. 4.**
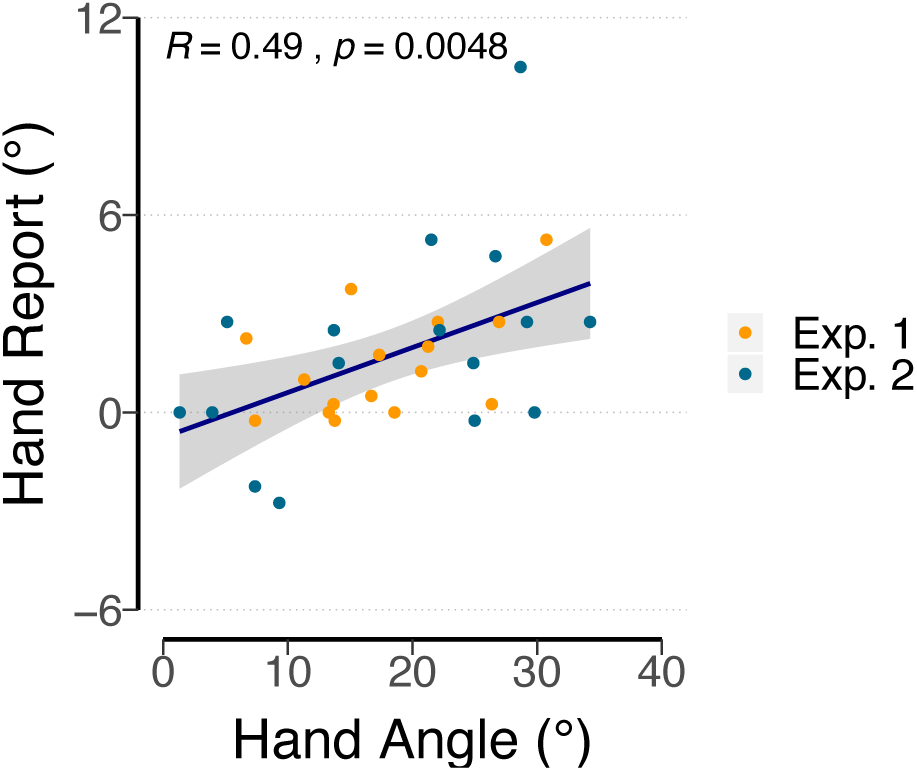
Pearson correlation between hand angle and hand reports during late adaptation, pooling together data from Experiment 1 (cycles 71 – 75, yellow dots) and Experiment 2 (end of first clamp block: cycles 66 – 70, blue dots). Correlations are marginally significant if data from each experiment were analyzed separately (Experiment 1: *R* = 0.5, *p* = 0.05; Experiment 2: *R* = 0.48, *p* = 0.059), likely reflecting a lack of statistical power. Solid line corresponds to the best-fitting regression line, while the grey shaded region corresponds to the 95% confidence interval.

Despite the contributions from proprioception, either associated with the veridical hand position or biases induced from the visual feedback (i.e., proprioceptive shift), the most striking feature of the data is that the hand reports remain close to the target location. This illusory experience likely reflects a third source of information underlying awareness during adaptation: The feedforward signal associated with a motor plan to reach to the target. Indeed, this signal appears to dominate both the veridical and distorted proprioceptive feedback (Izawa & Shadmehr, 2011; Ruttle, Hart, & Henriques, 2020), at least when the behavioral change is driven implicitly. To return to our opening example, the tired ping-pong player may be well aware of her state but is largely insensitive to the changes enacted by her brain to compensate for her fatigue.

## Author Contributions

All authors contributed to the study design. Testing, data collection, and data analysis were performed by J.S.T and D.E.P. All authors contributed to the interpretation of results under the supervision of R.B.I. J.S.T drafted the manuscript. D.E.P and R.B.I. provided critical revisions. All authors approved the final version of the manuscript for submission.

## Acknowledgements

We thank Odeya Kagan and Janet Hwang for their assistance with data collection. We thank Guy Avraham for astute comments on this manuscript. J.S.T was funded by a 2018 Florence P. Kendall Scholarship from the Foundation for Physical Therapy Research. This work was supported by grants NS092079 and R35NS116883 from the National Institutes of Health.

## Notes

### Competing Interest Statement

The authors have declared no competing interest.

